# Telomere DNA G-quadruplex folding within actively extending human telomerase

**DOI:** 10.1101/435545

**Authors:** Linnea I. Jansson, Joseph W. Parks, Jendrik Hentschel, Terren R. Chang, Rishika Baral, Clive R. Bagshaw, Michael D. Stone

**Affiliations:** Molecular, Cell and Developmental Biology Department, University of California Santa Cruz, CA 95060; Invitae, CA 94103.; Chemistry and Biochemistry Department, University of California Santa Cruz, CA 95060

**Keywords:** Telomerase, telomere, G-quadruplex, DNA structure, Ribonucleoprotein

## Abstract

Telomerase maintains telomere length by reverse transcribing short G-rich DNA repeat sequences from its internal RNA template. G-rich telomere DNA repeats readily fold into G-quadruplex (GQ) structures in vitro, and the presence of GQ-prone sequences throughout the genome introduces challenges to replication in vivo. Using a combination of ensemble and single-molecule telomerase assays we discovered that GQ folding of the nascent DNA product during processive addition of multiple telomere repeats modulates the kinetics of telomerase catalysis and dissociation. Telomerase reactions performed with telomere DNA primers of varying sequence or using K^+^ versus Li^+^ salts yield changes in DNA product profiles consistent with formation of GQ structure within the telomerase-DNA complex. Single-molecule FRET experiments reveal complex DNA structural dynamics during real-time catalysis, supporting the notion of nascent product folding within the active telomerase complex. To explain the observed distributions of telomere products, we fit telomerase time series data to a global kinetic model that converges to a unique set of rate constants describing each successive telomere repeat addition cycle. Our results highlight the potential influence of the intrinsic folding properties of telomere DNA during telomerase catalysis and provide a detailed characterization of GQ modulation of polymerase function.

**SIGNIFICANCE:** Telomeres protect the ends of linear chromosomes from illicit DNA processing events that can threaten genome stability. Telomere structure is built upon repetitive G-rich DNA repeat sequences that have the ability to fold into stable secondary structures called G-quadruplexes (GQs). In rapidly dividing cells, including the majority of human cancers, telomeres are maintained by the specialized telomerase enzyme. Thus, telomerase and its telomere DNA substrates represent important targets for developing novel cancer drugs. In this work, we provide evidence for GQ folding within the newly synthesized DNA product of an actively extending telomerase enzyme. Our results highlight the delicate interplay between the structural properties of telomere DNA and telomerase function.

## INTRODUCTION

Telomeres safeguard the ends of chromosomes from illicit DNA processing events that would otherwise threaten genome stability (1, 2). The foundation of telomere structure consists of short G-rich DNA sequence repeats. The majority of telomere DNA is double-stranded and can be up to several kilobases in length, while the ends are processed to terminate with a short 3’ single-stranded G-rich tail (∼50-150 nucleotides in vertebrates) (3, 4). Repetitive G-rich DNA sequences are not unique to telomeres and are found throughout the human genome (5). These G-rich repeats have the capacity to fold into G-quadruplexes (GQs), structures composed of multiple Hoogsteen bonded G-quartet motifs that stack together to yield stable DNA folds (6, 7). GQ folding has been implicated in a variety of biological processes. For example, replication of GQ-prone sequences is problematic and requires contributions from specific DNA helicase enzymes to avoid replication-coupled DNA damage (8–10). Sequences with GQ-folding potential are enriched within promoter sequences of oncogenes where they are thought to regulate gene expression (11). Finally, recent evidence suggests GQ folds can form in vivo in a spatially and temporally regulated manner (12–14). Thus, small molecules that bind and stabilize GQ-folds hold promise as novel cancer drugs; a fact that motivates better understanding of how GQ structure can modulate enzyme function.

Telomerase is an RNA-dependent DNA polymerase that is uniquely adapted to synthesizing G-rich repetitive DNA sequences (15, 16). Telomerase activity combats gradual telomere shortening that occurs with each round of cellular division (17). Left unchecked, telomere shortening induces senescence or cell death in somatic tissues. In contrast, proliferative cells such as stem cells rely upon telomerase activity to maintain telomeres in order to support continued rounds of cell division (15). Genetically inherited loss of function mutations in telomerase subunits cause human disorders characterized by deterioration of proliferative tissue types (18–21). On the other hand, telomerase overexpression contributes to the immortal phenotype of ∼90% of human cancers, and is therefore an important target for development of novel cancer therapies (22).

Telomerase is a ribonucleoprotein (RNP) complex that includes the long non-coding telomerase RNA (TR) and the catalytic telomerase reverse transcriptase (TERT) protein subunit (23, 24). In vertebrates, the 3’ end of TR includes canonical H/ACA box RNA motifs and associated proteins that promote TR stability and efficient telomerase RNP biogenesis (25). To initiate telomerase catalysis, the 3’ ssDNA telomeric tail base pairs with the TR template, forming a short RNA-DNA hybrid that is extended in the TERT active site (Fig. 1A). TERT utilizes a limited region of TR to direct synthesis of a defined GGTTAG hexameric telomere DNA repeat sequence (*k*_*pol*_) (Fig 1A). A unique property of telomerase is the ability to translocate on the DNA product (*k*_*trans*_) in order to recycle the integral TR template during processive addition of multiple telomere repeats prior to dissociation from the DNA product (*k_off_*) (Fig 1A)(26). This repeat addition processivity (RAP) implicitly requires multiple points of contact between telomerase and its DNA substrate, a notion that is consistent with data from a variety of telomerase systems identifying ‘anchor site’ DNA interactions outside the enzyme active site (27–30).

**Figure 1.**
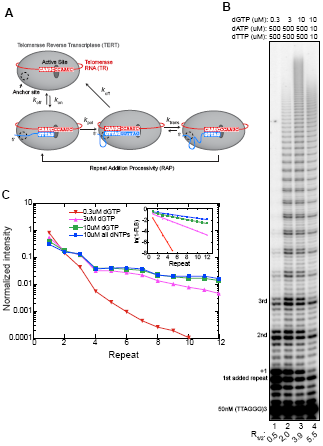
Human telomerase function. (A) Telomerase catalytic cycle. *k*_off_ and *k*_on_ represent the rate constants for dissociation and annealing to the telomere respectively. The rate constant for nucleotide addition during repeat synthesis is represented by *k*_pol_ and the translocation rate constant after the completion of each repeat is represented by *k*_trans_. The rate constants governing nucleotide addition and translocation together make up the Repeat Addition Processivity (RAP). TERT is represented by a grey oval and TR is shown simplified in red. The telomere primer is shown in blue and the telomerase anchor site is represented by a dashed circle. (B) Telomerase primer extension assay with 50mM ^32^P-labeled (TTAGGG)_3_ primer. Nucleotide concentration is indicated above the gel and repeats added to the (TTAGGG)_3_ primer are indicated on the left. The R_1/2_ value is shown at the bottom of the gel. (C) Normalized intensity plotted vs repeat number. Raw intensity for each band was converted to a fraction by dividing the band intensity by the total counts for the lane and then plotted against repeat number. The plot of ln(1-FLB) vs repeat number used to calculate processivity is inset in the top right corner.

Model telomere DNA substrates harboring integer multiples of four consecutive telomere repeats are inefficient binding substrates for telomerase in vitro while DNA primers with five, six, or seven consecutive repeats are efficiently bound and extended (31). Thus, while GQ structures can inhibit telomerase association, the presence of a small single-stranded DNA overhang in the substrate is sufficient to recover telomerase loading and function. These previous studies illuminated DNA sequence determinants that mediate the initial binding of telomerase to its substrate; however there remained an untested possibility that GQ structure may regulate the behavior of an actively extending telomerase-DNA complex, as suggested by studies of ciliate telomerase (32, 33). Furthermore, we reasoned that the specialized telomerase system provides a powerful opportunity to investigate the influence of GQ-forming sequences on nucleic acid polymerase function.

To study the relationship between DNA structure and human telomerase catalysis we performed direct primer extension assays using dNTP concentrations similar to those found in the cellular environment (34). Our experiments reveal a complex pattern of telomerase DNA product accumulation that indicates the efficiency of template recycling is dependent upon the number of synthesized repeats. Experiments using telomere DNA primers of varying sequence and varying salt conditions support the notion that a GQ can form within the telomerase-DNA complex. Single-molecule FRET experiments provide further support for DNA structural dynamics within actively extending telomerase enzymes. To estimate individual rate constants for successive repeat addition cycles we performed global kinetic modeling of telomerase time-series data. Interestingly, our model converges to a unique solution of rate constants that provides a direct measure of processivities for each cycle of telomere repeat addition. These results are consistent telomere DNA GQ folding serving to promote template recycling, as well as to accelerate product dissociation. We present a mechanistic model that provides a framework to understand the delicate interplay of telomere DNA structure and function during telomerase catalysis.

## RESULTS

### Telomerase product distribution is sensitive to dNTP concentrations and stoichiometry

When measuring telomerase activity in vitro, it is common to employ direct primer extension assays in the presence of α^32^P-dGTP. This approach permits reactions to be performed with a large excess of unlabeled DNA substrate, benefits from very high sensitivity of product detection, and circumvents PCR-induced artifacts inherent to the telomere repeat amplification protocol (TRAP) assay. However, the use of α^32^P-dGTP incorporation to detect product accumulation limits the amount of total dGTP that can be used in the assay, leading to the widely reported practice of using non-physiological dNTP stoichiometry that has the potential to significantly alter the telomerase product distribution (35, 36). To circumvent this problem, we used 5’ radiolabeled DNA primers and cold dNTPs to monitor telomerase activity (Fig. 1B). We compared this approach to standard assays performed with α^32^P-dGTP using identical dNTP and primer concentrations (Fig. S1). Although the reaction profiles are qualitatively distinct, we observed quantitatively similar product distributions for the two approaches when normalized for the amount of α^32^P-dGTP incorporation. Importantly, the majority of the input DNA primers are not extended in our experiments, demonstrating our reaction conditions are sufficient to limit distributive telomerase activity (i.e. an individual primer being extended by multiple telomerase enzymes). Processive telomerase activity is also evident when analyzing pulse-chase experiments in which longer DNA products continue to accumulate after addition of a 400-fold excess of cold DNA primer (Fig. S2).

**Figure 2.**
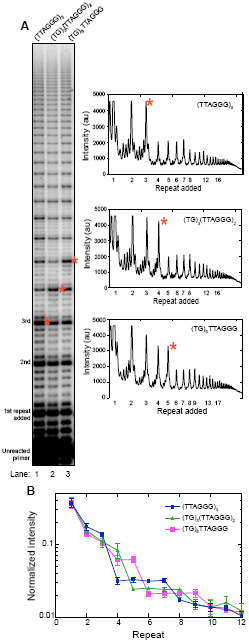
Product distribution profile is dependent on amount of consecutive TTAGGG repeats. (A) Telomerase primer extension assay with primers of varying TTAGGG composition. Primer variants are indicated at the top of the gel. Repeats added to the primer are indicated to the left. Lane profiles with raw intensity versus repeat band for each primer variant are shown on the right. Corresponding bands between the gel and lane profiles are indicated by a red asterisk. (B) Normalized intensity plotted vs repeat number. Fractional intensity of each band compared to total lane counts is plotted against repeat number. Data are averaged from three independent experiments.

Having established that our end-labeled DNA primer assay is capable of accurately monitoring processive telomerase action, we next sought to analyze the influence of varying dNTP concentrations on the telomerase product distribution (Fig. 1B). Previously published studies defined telomerase processivity as the number of repeats corresponding to the point where the dissociated DNA represents 50% of the total population (R_1/2_) (37, 38). This approach is analogous to half-time (t_1/2_) analysis and can be performed by fitting a linear regression to a plot of ln(1-FLB) versus repeat number, where FLB is the fraction left behind (Fig. 1C inset, see Methods for details). Titrating increasing amounts of dGTP in the presence of a large excess of dATP and dTTP yields a significant boost in RAP as has been reported previously for both human and *Tetrahymena* telomerase (Fig. 1B and 1C) (35, 39–43). However, the use of a large excess of dATP and dTTP is not a good approximation for the physiological dNTP pool which is generally closer to the ∼10 uM range (34). Note that dCTP is not required for the human telomere sequence and its absence does not alter telomerase function (data not shown). When assayed in the presence of equimolar dGTP, dATP, and dTTP, we observe the highest RAP activity of all conditions tested (Fig. 1B, lane 4 and Fig 1C). Moreover, using this approach, we noticed that the previously observed doublet band at the +1 position after each complete telomere repeat is effectively suppressed (Fig. 1B, compare lanes 1-3 with lane 4). We interpret this +1 band to reflect promiscuous use of the non-template RNA base at hTR position U45, which may be enhanced by the large excess of dATP. Based upon these results, we elected to perform all subsequent telomerase assays in our study under conditions of equimolar dNTPs that better reflect cellular dNTP concentrations.

### G-quadruplex folding alters the pattern of telomerase product accumulation

Established methods for approximating RAP using the R_1/2_ value described above assume an exponential decay in the distribution of accumulated telomerase product lengths with each telomere repeat added (37, 38, 44). However, we noted the appearance of plateaus in the product distribution when using equimolar concentrations of dNTPs (Fig. 1C and 2A). For example, when using a standard telomere DNA primer composed of the sequence (TTAGGG)_3_, we observe a sudden drop in product accumulation between the bands corresponding to the third and fourth telomere repeats added to the primer (Fig. 2A, red asterisk). Surprisingly, the intensities of the subsequent four added repeats are approximately equal, until a second decrease in accumulation occurs between added repeats seven and eight. This pattern of four equally populated product lengths, followed by a decrease in accumulation, continues throughout the detectable range of telomere DNA products.

Telomere DNA primers with at least four contiguous G-rich repeats can fold into a GQ in vitro (45, 46), suggesting the RAP-associated ‘pattern of four’ we observe in our experiments may be due to GQ folding of the DNA product within an actively extending telomerase complex. To test this hypothesis, we altered the 5’ end of the telomere DNA sequence so that it no longer harbored the requisite run of guanines needed to participate in GQ folding (Fig. 2A, lanes 2 and 3). Altering the primer in this way should change the product length where the ‘pattern of four’ appears if the newly synthesized DNA folds into a GQ. Indeed, a modified DNA primer with a 5’ (TG)_3_ substitution supports telomerase RAP, but the plateaus in the product profile are delayed by one additional repeat, corresponding to the sequence needed to promote GQ formation in the product DNA (Fig. 2A, compare lanes 1 and 2). Similarly, when the first two repeats in the telomere DNA primer were substituted with the TG dinucleotide sequence ((TG)_6_), the plateaus are delayed by two additional repeats (Fig. 2A, compare lanes 1 and 3). These results were highly reproducible across three independent experimental trials (Fig. 2B) and agree with the hypothesis that GQ folding within the nascent telomere DNA causes the telomerase product profile to deviate from a strictly decreasing decay.

### GQ stabilization alters telomerase kinetics and DNA product structural dynamics

The H-bonding configuration of the G-quartet motifs within a GQ fold can be differentially stabilized by coordination of specific monovalent cations, with a rank order of K^+^ > Na^+^ > Li^+^ in terms of degree of stabilization (47). Therefore, we next set out to further dissect the contribution of GQ folding to telomerase activity by altering the salt conditions of our telomerase assays. We observed robust telomerase activity in all salt conditions tested; however, there was a clear reduction in total product accumulation in Li^+^ when compared to Na^+^ and K^+^ (Fig. 3A). The lower total product accumulation in the presence of Li^+^ is a consequence of slower DNA synthesis kinetics (Fig. S3) and is not completely unexpected based on reported effects of Li^+^ on DNA polymerases (48). Interestingly, we do not observe the robust “pattern of four” RAP product distribution in the presence of Li^+^ (Fig. 3A and 3B), the salt condition expected to least stabilize GQ folding during telomerase catalysis. This result lends additional support to the hypothesis that GQ formation in the presence of K^+^ or Na^+^ impacts the product distribution of an actively extending telomerase complex.

**Figure 3.**
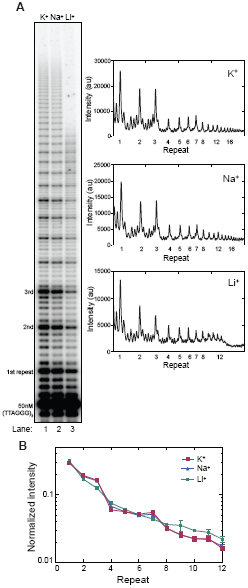
Product distribution profile is dependent on monovalent salt identity. (A) Telomerase primer extension assay in different salt conditions. Repeats added to the primer are indicated to the left. Lane profiles with raw intensity versus repeat band for each lane are shown on the right. (B) Normalized intensity plotted versus repeat number. Fractional intensity of each band compared to total lane counts is plotted against repeat number. Data are averaged from three independent experiments.

The results of our ensemble telomerase assays suggest that folding of the nascent DNA product can influence telomerase catalysis. However, this approach does not directly interrogate DNA conformation within an active RNP complex. Therefore, we turned to a single molecule Förster Resonance Energy Transfer (smFRET) based approach that directly monitors DNA structure and dynamics within individual telomerase enzymes (49, 50). In order to ensure our smFRET assay supports telomerase activity in both K^+^ and Li^+^ buffers we employed a recently reported method for in situ detection of extended DNA products at the single-molecule level (51). Telomerase RNP complexes harboring a Cy3 dye incorporated into the telomerase RNA subunit were bound to a biotinylated DNA primer and then surface-immobilized onto a streptavidin coated glass slide (Fig. 4A). The telomerase-DNA complexes were incubated in either K^+^ or Li^+^ activity buffer as well as with a Cy5-labeled detection oligonucleotide with a sequence that is complementary to the telomere product. In this way, telomere primers that are being actively extended by telomerase (i.e. enzyme is still bound and some of the DNA product becomes accessible) are detected as a FRET signal between the Cy3-labeled enzyme and the Cy5-labeled DNA probe. The appearance of the FRET signal was strictly dependent upon addition of activity buffer containing dNTPs and was time dependent (Fig. 4B). After 20 minutes of incubation, comparable levels of telomerase activity, measured as the total number of telomerase-DNA complexes producing a positive FRET signal, were detected in both K^+^ and Li^+^ activity buffers (Fig. 4B). Taken together, these results demonstrate that telomerase is catalytically active in both K^+^ and Li^+^ buffers in the single-molecule assay.

**Figure 4.**
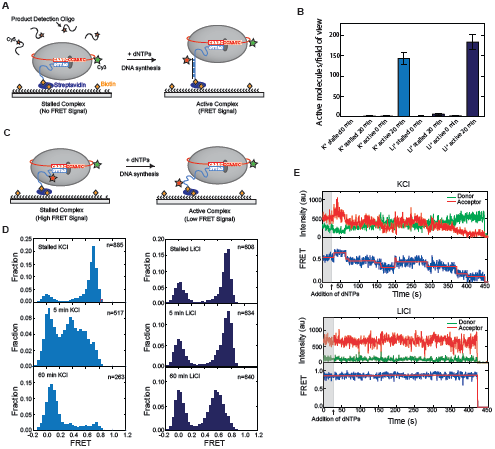
smFRET studies of telomerase in the presence of K ^+^ and Li^+^. (A) Schematic of human telomerase smFRET activity assay. Purified telomerase is immobilized to a pegylated and biotinylated quartz slide through binding to a biotinylated telomere primer (blue). TERT is depicted as a grey oval and hTR is shown in red. A Cy3 dye (green star) is conjugated to hTR. A Cy5-labeled detection oligo complementary to two and a half telomere repeats is added to stalled enzyme (left). Upon addition of dNTPs, telomere synthesis begins, and sufficient lengths of telomere DNA for the Cy5-detection oligo to anneal to are extruded from the telomerase active site (right). (B) Telomerase is active in both K^+^ and Li^+^. smFRET experiments demonstrate a significant increase in active molecules per field of view in both KCl and LiCl buffers only in the presence of dNTPs. Experiments performed in KCl are shown in light blue and experiments performed in LiCl are shown in dark blue. The number of active molecules per field of view was calculated from the number of FRET-positive spots, as an indicator for the presence of Cy5-probe that annealed to actively synthesized telomere repeats. (C) Schematic of human telomerase smFRET experimental designed to probe DNA conformational change. The telomere primer (blue) is conjugated to a Cy5 dye (red star). Upon addition of dNTPs, telomere repeat synthesis will move the Cy5 dye on the telomere primer further from the active site, resulting in a lower FRET value (right). (D) smFRET histograms in K^+^ and Li^+^. Histograms depict the FRET distribution for stalled telomerase (top panels) as well as after 5 and 60 minutes after addition of dNTPs (middle and bottom panels respectively). Experiments performed in KCl are shown in the left panels and experiments performed in LiCl are shown on the right. (E) Real time smFRET traces. Representative FRET traces are shown for experiments performed in KCl (top panel) and LiCl (bottom panel). Donor intensity is shown in red and acceptor intensity in green. The corresponding FRET value (blue) was fit with steps (red) using MATLAB.

Having detected robust telomerase activity in both K^+^ and Li^+^ buffers, we next performed smFRET assays with Cy3-labeled telomerase and Cy5-labeled telomere primer in order to directly monitor DNA dynamics during active telomere elongation. (Fig. 4C). Stalled telomerase-DNA complexes harboring these site-specific dye modifications were surface-immobilized as described above. Data collected on stalled complexes prepared in this manner yielded a unimodal FRET distribution centered at ∼0.75 (Fig. 4D, top left panel). Next, the telomerase complexes were activated for DNA synthesis by introducing dNTPs in telomerase activity buffer (containing 50mM KCl), resulting in a significant shift towards lower FRET values and ultimately yielding a single population centered at near-zero FRET (Fig. 4D, left panels). This FRET change upon activation of DNA synthesis is consistent with the Cy5-label site on the telomere DNA moving further away from the telomerase active site, and perhaps folding into a stable structure, during multiple rounds of DNA repeat synthesis. Analysis of real-time single-molecule FRET trajectories collected on actively extending telomerase complexes in this same condition reveals complex DNA conformational dynamics characterized by a general transition to discrete lower FRET states combined with the occurrence of transient increases in FRET (Fig. 4E and Fig. S4).

In order to dissect the contribution of DNA structure to the observed FRET dynamics we performed the same experiment using LiCl instead of KCl in the telomerase activity buffer. Once again, the initial FRET population of stalled telomerase-DNA complexes was centered at ∼0.75 in the presence of Li^+^ (Fig. 4D, top right panel). However, in contrast to the K^+^ activity buffer, we did not observe a substantial drop in FRET upon addition of dNTPs (Fig. 4C right panels). Instead we observed a modest shift in the major FRET population to ∼0.6 after 60 minutes incubation time. This result is consistent with individual smFRET trajectories of telomerase-DNA complexes incubated in Li^+^ activity buffer, where the FRET level appears to remain constant over the course of several minutes (Fig. 4E). Together, these results argue that the formation of a GQ in the nascent telomere product contributes to efficient extrusion of the telomere DNA away from the telomerase active site during the course of activity.

### Global kinetic modeling provides a measure of telomerase microscopic processivity

Telomerase processivity can be modeled as a series of consecutive reactions in which nucleotide addition is in competition with DNA dissociation at each step of the reaction. To simplify our telomerase kinetics analysis, we focus on the intense repeat addition bands, assuming the intervening nucleotide addition steps are rapid and accompanied by little DNA dissociation. The macroscopic processivity of telomerase can be conveniently described by the median product length (R_1/2_= ln2/decay constant) (see Fig. 1C) (37). However, treating the data in this manner has several limitations. First, in a typical primer extension experiment one cannot distinguish between dissociated product and DNA that remained associated with enzyme at the point when the reaction was arrested. In addition, this analysis approach has the effect of suppressing possible deviations in microscopic processivity, which reflects the individual probability of adding another DNA repeat at a specific step of telomerase catalysis. The experiments described in the present study provide clear evidence that telomerase products do not accumulate uniformly and display patterns dependent upon assay conditions, DNA sequence, and/or product length. We therefore developed a kinetic model that can be utilized to globally fit telomerase time-series data in order to extract microscopic processivity values that underlie the observed distributions of telomerase products (Fig. 5A). Using this scheme, we treat this multistep process as a first-order reaction with an effective forward rate constant (*k_f_*) for the transition between each repeat and a dissociation rate constant (*k_d_*) (Fig. 5A) (note that *k*_*f*_ in this model reflects a combination of *k*_pol_ and *k*_trans_ described in Fig. 1A). Using the kinetic scheme depicted in Fig. 5A, we can then define the microscopic processivity (p) at each step as p = *k_f_* / (*k_f_* + *k*_*d*_).

**Figure 5.**
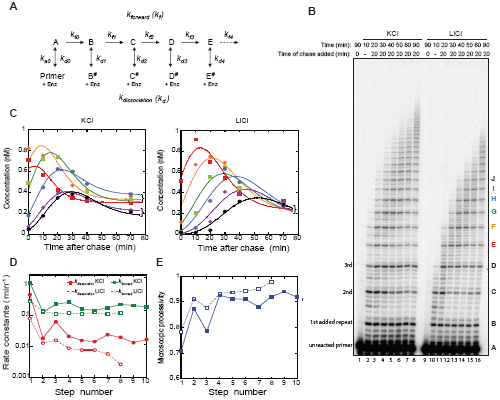
Human telomerase kinetics. (A) Kinetic mechanism for processive telomerase activity used to globally fit the primer extension assay shown in (B). The letters refer to the repeat band number (B = 1st added repeat, C = 2nd repeat etc.) and dissociated products are identified with the # symbol. Band intensities are proportional to the sum of products (e.g. B + B^#^). (B) Extending primer dissociation rate assay in the presence of KCl and LiCl. Primer extension assays were performed with 50 nM ^32^P-labeled (TTAGGG)_3_ primer. 20 uM cold (TTAGGG)_3_ primer was added to the reaction after 20 minutes of activity. A control reaction with 20uM cold primer added at the beginning of telomerase activity was included for both buffer conditions (lanes 1 and 9). Repeat number added to the primer is indicated on the left of the gel. Letters indicating band identity for kinetics modeling is indicated on the right of the gel. Bands EJ are colored according to plot shown in (C). (C) Representative global fits to bands E-J in KCl (left panel) and LiCl (right panel). The concentration of the products (see color code in (B)), based on band intensity relative to the initial 50 nM primer, was plotted against the time after the cold chase. Note the clustering of bands E-H and I-J 70 minutes post chase in the presence of KCl, which corresponds to the four repeats of the first plateau and the first two repeats of the second plateau (cf. Fig. 2). This partitioning is not present in the presence of LiCl. (D) Consecutive rate constant values for forward repeat addition (green squares, *k*_*f*_) and dissociation (red circles, *k*_*d*_) returned by DynaFit for data in the presence of KCl (solid symbols) and LiCl (open symbols). The step number refers to the rate constant subscripts shown in (A). Note the overall reaction is slower in the presence of LiCl and beyond band I (8th step) the fitted rate constant values had a large error because the decay phase had barely started by 70 min. Therefore, these values were omitted. (E) Microscopic processivity (*k*_*f*_ / (*k*_*f*_ + *k*_*d*_)) at each step of the reaction calculated from the rate constants shown in (D). Note the saw-tooth structure in the presence of KCl (solid line) compared to the relative lack of structure in LiCl beyond the second step (dashed line).

DNA dissociation is an effectively irreversible process when a large excess of unreacted primer remains, which outcompetes the re-binding of any product DNA. To ensure our experiments complied with this assumption, we analyzed telomerase kinetics following a chase with 400-fold excess of unlabeled primer DNA, which serves to block re-association of the labeled DNA primer following telomerase dissociation (Fig. 5B). Telomerase time course assays were performed in the presence of either K^+^ or Li^+^ activity buffer conditions (Fig. 5B). Activity was initiated at time zero in the presence of end-labeled telomere DNA primer and dNTPs, followed by addition of excess chase primer at 20 minutes. The presence of 400-fold excess cold primer prior to enzyme addition was sufficient to eliminate any observable extension of the 50 nM endlabeled DNA primer used in our assays (Fig. 5B, lanes 1 and 9). Time points were collected at regular intervals out to 90 minutes and the concentration of each repeat species (B + B^#^, C + C^#^, etc, Fig 5A) was determined at each time point from the band intensity, knowing that the intensity of the initial primer was 50 nM. Individual rate constants were estimated by fitting the concentration time courses globally, using DynaFit (Fig. 5C, data shown for bands E-J). Global fitting converged to a unique set of rate constants that could be extracted with reasonable precision (see Supplementary Methods for details of kinetic modeling) (Fig. 5D). Comparison of the data obtained in the presence of K^+^ and Li^+^ revealed that the rate constants, *k_f_* and *k_d_*, decrease with increasing repeat number and result in lower microscopic processivity values for short DNA products as has been noted previously (37). However, in the presence of K^+^, the rate constants and microscopic processivity values show a saw-tooth modulation that gives rise to the ‘pattern of four’ clustering of products noted earlier (Fig. 5E, closed symbols). Although the effect is relatively small it is robust as determined by Monte-Carlo analysis and is reproducible between experiments and telomerase preparations (Fig. S5 and S6). Interestingly, the rate constants for repeat addition (*k_f_*) and dissociation (*k_d_*) were greater in the presence of K^+^ than in Li^+^, but the resultant microscopic processivity was lower (Fig. 5D and 5E). Taken together, these results suggest that in the presence of K^+^, the ability of the DNA product to form a GQ fold (which first arises at step 3, see Discussion for details) appears to promote translocation (increased *k_f_*) but at the increased risk of DNA product dissociation (increased *k*_*d*_).

## DISCUSSION

The foundation of telomere structure consists of short G-rich repeat sequences, GGTTAG in humans, that have the propensity to fold into G-quadruplex (GQ) structures in vitro and in vivo (6, 7). Short ssDNA oligonucleotides that form GQs are poor telomerase substrates in vitro, due to the occlusion of an accessible single-stranded 3’ end (31, 46). Here, we present evidence that GQ formation can occur within an actively extending telomerase complex in vitro and that formation of such GQs affect the kinetic properties of telomerase. Telomerase activity assays using modified telomere DNA primer sequence and varying salt conditions together support the notion that GQ folding occurs during telomere DNA synthesis. Consistent with this concept, single-molecule FRET experiments directly demonstrate complex folding dynamics of the nascent DNA product in the presence of K^+^ but not Li^+^. We describe a detailed kinetic framework for telomerase catalysis and use this model to globally fit telomerase time-series data in order to extract microscopic processivity values for each cycle of telomere DNA repeat synthesis (see below for details). Our kinetic modeling reveals small but significant GQ-dependent changes in the rate constants describing the DNA repeat synthesis reaction (*k_f_*) and product dissociation (*k*_*d*_).

Telomerase repeat addition processivity (RAP) can be quantitatively described as the number of repeats corresponding to the point where the dissociated DNA represents 50% of the total population (R_1/2_) (37). The median number of repeats is a measure of the macroscopic processivity of telomerase that is implicitly assumed to be independent of product length. This method of analysis has been useful in previous studies that sought to characterize telomerase RAP under varying experimental conditions or with mutant telomerase enzymes (37, 52). A caveat of this approach is that typical telomerase primer extension assays cannot distinguish between products that have already dissociated and DNA that remained bound to enzyme when the reaction was stopped. This caveat can be addressed by the kinetic modeling approach described in the present study, or by introducing additional experimental steps to separate dissociated and bound products prior to gel analysis (43). However, one additional advantage of the global kinetic modeling approach is the ability to extract microscopic processivity values that reflect the individual probability of synthesizing each subsequent telomere repeat. Indeed, the results of our global kinetic modeling reveal a saw-tooth modulation of *k_f_* and *k_d_*, that together explain the ‘pattern of four’ clustering of telomerase product accumulation.

We present a working model for the mechanism of GQ-dependent effects on telomerase repeat addition processivity (Fig. 6). The complex rearrangements that are necessary for template recycling during multiple rounds of telomere repeat synthesis require multiple points of contact between telomerase and its DNA substrate. Minimally, the DNA must engage the enzyme by interacting with the RNA template in the active site and/or making distal contacts with a separate ‘anchor site’ (27–30). Upon completion of a telomere repeat, the 3’ end of the DNA product must dissociate from the template and dynamically realign with the downstream region of the RNA to prime the next round of repeat synthesis (Fig. 6) (50). This large-scale translocation step represents a vulnerable stage of telomerase catalysis, requiring a stable anchor-site interaction to prevent product dissociation. In principle, GQ folding within the DNA product can compete with the anchor-site contact to promote dissociation (Fig. 6, bottom pathway). Alternately, GQ folding can bias the positioning of the 3’ end of the primer to favor realignment for a subsequent round of repeat synthesis while allowing for anchor site contacts to remain intact (Fig. 6, top pathway). Which pathway the enzyme takes will depend on if anchor site interactions are maintained and the register of the 3’ end of the telomere during the formation of a GQ. In the schematic model depicted in Fig. 6, a GQ formed when the most recently synthesized telomere is bound in the active site will disrupt anchor site interactions and promote product dissociation, while a GQ formed when the 3’ telomere end is annealed to the template priming site, will facilitate another round of repeat addition. The latter outcome is mechanistically similar to DNA hairpin induced translocation models proposed for diverse telomerase systems (53, 54).

**Figure 6.**
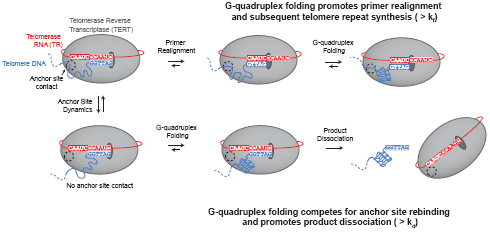
Model of GQ folding within actively extending telomerase complex. At the completion of a telomere repeat, telomerase (with TERT depicted in grey and hTR in red), is annealed to the telomere primer (blue). The anchor site (dashed circle) is either engaged with the telomere DNA (upper pathway) or disengaged (lower pathway). If the anchor site contacts are maintained and primer realignment occurs (upper, middle carton), there should be sufficient DNA to allow for a GQ to form within the actively extending enzyme (top, right cartoon). The formation of a GQ at this stage may bias the enzyme complex towards another round of telomere repeat addition. If the anchor site contacts are broken (lower, left cartoon) while the 3’ end of the telomere is bound in the active site, there may be competition between GQ formation and anchor site interactions (bottom left and middle cartoons). If primer realignment occurs at this stage, product dissociation will occur (bottom, right cartoon).

The finding of GQ-dependent effects on telomerase catalysis in vitro begs the question of whether a similar phenomenon occurs in the cell. While direct demonstration of GQ folding in vivo is challenging, recent experimental advances using GQ specific antibodies suggest that GQ folding can occur in a temporally and spatially regulated manner in vivo (12–14). Additionally, studies of ciliate telomerase have suggested that GQ folding may contribute to product dissociation during active telomere elongation (32, 33). The recently reported cryo-EM structures of the human and *Tetrahymena* telomerase enzymes provide a platform for investigating the potential for GQ folding within the actively extending telomerase complex (55, 56). Interestingly, when analyzing the EM density of the human telomerase complex, we noticed a structural pocket immediately proximal to the path of the nascent DNA emerging from the active site (Fig. S7). This pocket is flanked by two evolutionarily conserved features that are essential for telomerase processivity: the TEN domain and the TR p6.1 stem loop. The volume of this pocket is remarkably compatible with the dimensions of a single GQ fold, and raises the possibility that the nascent DNA products has ample space to assume a GQ structure within the confines of telomerase RNP complex. In the *Tetrahymena* structure, the C-terminal domain of the processivity factor, Teb1, is positioned as a lid to the DNA exit pocket, forming an enclosed cavity that is also sufficient to accommodate a single GQ fold. We propose the hypothesis that the Teb1-related shelterin component POT1 may occupy a similar position in the human telomerase complex. Thus, the anticipated binding of POT1 to the DNA as it is threaded out of this DNA exit channel is not mutually exclusive with the possibility of GQ folding within this protected cavity (46, 56). While speculative, the notion of GQ folding within the DNA exit channel is consistent with the amount of DNA protected by exonuclease cleavage as well as previous smFRET measurements that showed a similar amount of the DNA substrate is in close proximity to the template (50, 57). Future structural and biochemical experiments are required to investigate the influence of the shelterin proteins POT1-TPP1 on the folding properties of the nascent telomere DNA product, as well as the in vivo significance of GQ folding during telomerase catalysis.

## MATERIALS AND METHODS

### Preparation of RNAs

#### Synthetic, dye-labeled RNA fragments

Synthetic PK hTR fragment 32-62 was ordered from Dharmacon with an internal aminoallyl uridine (5-N-U) on position U42. A single tube of RNA was resuspended in nuclease-free H2O and ethanol precipitated in the presence of 300mM NaOAc pH 5.2 to prevent introduction of reactive amines. The pellet was resuspended in 100 ul 0.1M NaHCO3 and was used to solubilize a single mono-reactive Cy3-dye pack (Amersham). The solution was incubated for 2 h at 37°C. The solution was then ethanol precipitated, again with NaOAc. The RNA was then deprotected by resuspension in deprotection buffer (100mM acetic acid, pH 3.6) followed by vortexing and centrifugation for 10 seconds each. The solution was then heated at 60°C for 30 min followed by ethanol precipitation. The pellet was resuspended in 60 ul 0.1M TEAA, pH 7.5, and was HPLC purified on a reverse phase C18 column (Agilent technologies).

#### Ligation of RNA fragments

A splinted ligation reaction containing 800 pmol of Cy3-labeled hTR 32-62 fragment, 1600 pmol of in vitro transcribed unlabeled hTR 63-195, 1600 pmol DNA splint and 0.5x ligase buffer (NEB) was set up in a 200ul volume and incubated for 95C for 5 min followed by 30C for 10 min. 200ul ligation mix (1.5x ligase buffer, 8000 units of T4 DNA ligase (NEB), 2 mM ATP and 1U/ul RNasin Plus) was then added to the reaction mixture and incubated overnight at 30°C. 10 units of Turbo DNase was added and incubated for 15min at 37°C. The RNA was then phenol-chloroform extracted and ethanol precipitated prior to PAGE purification.

#### In vitro transcription

Unlabeled CR4/5 (hTR 239-328) and PK (32-195) was in vitro transcribed using homemade T7 polymerase in 1x RNA polymerase buffer (40mM Tris-HCl pH 7.9, 6mM MgCl_2_, 1mM DTT, 2mM spermidine) with 1.5 mM NTPs, 22 mM additional MgCl_2_, 90mM additional DTT and 40 units RNasin Plus. The reaction was incubated overnight at 37°C followed by the addition of 10 units of Turbo DNase for 15min at 37C. The RNA was phenol-chloroform extracted and ethanol precipitated prior to PAGE purification. PK 63-195 was in vitro transcribed as above but with 1mM of each NTP and an additional 5mM GMP in order to preferentially make a 5’ monophosphate fragment for use in splinted ligation.

### Telomerase expression

Human telomerase was reconstituted using the Promega TnT Quick Coupled

Transcription/Translation system. In Lo-bind tubes (Eppendorf), 200ul of TnT quick mix was combined with 5ug of pNFLAG hTERT plasmid as well as 1uM of the in vitro transcribed PK and CR4/5 fragments except for when using Cy3-labeled PK, which was added at 0.1uM. The reaction was incubated for 3 h at 30°C. 5 ul of 0.5 M EDTA, pH 8.0, was added to quench excess Mg^2+^ present in the lysate.

### Telomerase purification

Directly after reconstitution, telomerase was purified using the N-terminal FLAG tag on hTERT. To pull down the enzyme, Sigma Anti-FLAG M2-agarose resin was used. 50 ul bead slurry was first washed three times with wash buffer (50 mM Tris-HCl, pH 8.3, 3 mM MgCl_2_, 2 mM DTT, 100 mM NaCl) with 30 sec spins at 5000rpm at 4°C after each wash. The beads were then blocked in blocking buffer (50mM Tris-HCl pH 8.3, 3mM MgCl_2_, 2mM DTT, 500ug/ml BSA, 50ug/ml glycogen, 100ug/ml yeast tRNA) for 15 min under gentle agitation at 4°C followed by a 30 sec spin and removal of the supernatant. This blocking step was performed twice. After blocking, the beads were resuspended in 200ul fresh blocking buffer and added to the telomerase lysate. The beads and lysate were incubated for 2 h at 4°C under gentle agitation. The beads were then spun down for 30 sec at 5000 rpm and at 4°C and the supernatant was discarded. The beads were then washed three times in wash buffer containing 300mM NaCl (KCl or LiCl for salt dependence experiments) followed by three washes in wash buffer with 100 mM NaCl (KCl or LiCl for salt dependence experiments). A 30 sec spin at 5000 rpm at 4°C was performed between each wash. To elute the enzyme, the beads were incubated in 60 ul elution buffer (50 mM Tris-HCl, pH 8.3, 3 mM MgCl_2_, 2 mM DTT, 750 ug/ml 3x FLAG peptide, 20% glycerol) under gentle agitation at 4°C for 1 h. After elution, the beads were removed by centrifugation at 13000 rpm through Nanosep MF 0.45um filters. 5ul aliquots were then aliquoted into Lo-bind tubes and flash frozen in liquid nitrogen and stored at −70°C until use.

### End-labeling of primers

50pmol of primer was labeled with alpha-^32^P ATP using T4 PNK in 1x PNK buffer (70 mM Tris-HCl, pH 7.6, 10 mM MgCl_2_, 5 mM DTT) in 50 ul. The reaction was incubated for 1h at 37°C followed by heat inactivation of T4 PNK at 65°C for 20min. Centrispin columns (Princeton separations) were used to obtain labeled primer.

### Primer extension assays

Telomerase activity assays were performed by using 5 ul of purified telomerase in 1x activity buffer (50 mM Tris-HCl pH 8.3, 50 mM KCl (NaCl or LiCl when indicated), 1 mM MgCl_2_, 2 mM DTT) in a final volume of 15ul. For experiments containing end-labeled primer, each reaction contained 10 uM each of dATP, dTTP and dGTP as well as 50 nM of indicated ^32^P-primer. For reactions containing radiolabeled dGTP, each reaction contained the indicated dNTP concentrations as well as 50nM unlabeled (TTAGGG)_3_ primer. Reactions were incubated for 90 minutes at 30C. Reactions were stopped using 200ul 1x TES buffer (10 mM Tris-HCl, pH 7.5, 1 mM EDTA, 0.1% SDS), followed by phenol-chloroform extraction and ethanol precipitation of the DNA. The pellet was resuspended in 1x formamide gel loading buffer and resolved on a 12% denaturing PAGE gel. The gel was then dried and exposed on a phoshorimager screen and scanned using a Typhoon scanner. Band intensities were quantified using SAFA (58). R1/2 values were calculated by summing each band and all bands below it divided by the total counts for a given lane (to calculate the fraction left behind (flb). The natural log of (1- flb) was then plotted against repeat number and 0.693 was divided by the value of the slope to give the R1/2 value (37).

### Single-molecule experiments

#### Slide preparation

Quartz slides (Finkenbeiner Inc.) were boiled in water for 20 min to remove parafilm, epoxy and coverslips from previous experiments. The quartz slides were then scrubbed with alconox by hand and rinsed with MilliQ water. The slides were then placed in a solution containing 10% w/v Alconox and sonicated for 20 min. The slides were then rinsed and sonicated in MilliQ water for 5 min. The slides were then sonicated in acetone for 15 min followed by direct transfer to a solution containing 1 M KOH and sonicated for 20 min. The slides were then rinsed with water and thoroughly flame dried using a butane torch (BernzOmatic). After flaming, the slides were left to cool on a drying rack in the fume hood. While the slides were cooling, a slide holder containing methanol was sonicated for 5 min and a silanizing solution containing 100 ml methanol, 5 ml of glacial acetic acid and 1 ml of N-(2-aminoethyl)-3- aminopropyltrimethoxysilane (UCT) was prepared during this time. After the slides had cooled down, the methanol in the slide holder was exchanged for the silanizing solution and the slides were sonicated in this solution for 1 min and then allowed to stand in the solution for an additional 20 min at room temperature. While slides were incubating, 400 mg of mPEG-Succinimidyl Valerate MW 5000 (Laysan Bio, Inc.) was resuspended in 800 ul 0.1M NaHCO_3_. Also, 2 mg of Biotin-PEG-Succinimidyl Valerate MW 5000 (Laysan Bio, Inc.) was resuspended in 200 ul of 0.1M NaHCO_3_ and mixed with the other PEG solution. After combining, the PEG solution was briefly sonicated to ensure complete resuspension. The slides were rinsed with MilliQ water and dried with nitrogen gas and then put in a humidor box. 150 ul of the PEG solution was applied to each slide and covered with a coverslip. The slides were incubated in the dark overnight. The next day the coverslip and PEG solution was rinsed off using MilliQ water and dried using nitrogen gas. Channels were assembled using parafilm strips as spacers on the pegylated quartz slide and plasma-cleaned coverslips were added as the upper channel face. For real-time experiments, flow cells were assembled by first drilling two holes at opposite ends of a quartz slide. A cut pipette tip was glued into one hole to act as a reservoir, and intramedic® PE100 polyethylene tubing glued to the other to provide a vacuum for buffer exchange. The channel was cut from a piece of double sided tape which was sandwiched between the pegylated quartz slide and a coverslip.

#### Enzyme immobilization

Channels were incubated with 30 ul 10 mg/ml BSA (NEB) for at least 30 min. 60 ul of 0.2 mg/ml MPS (streptavidin) was then added and incubated for 5 min. The channel was then washed with 150 ul T50 buffer (10 mM Tris-HCl, pH 8, 50 mM NaCl, or LiCl for Li^+^ experiments)). 2.5 ul of Cy3-telomerase, 0.5 nM primer-Cy5 and Ix imaging buffer (50 mM Tris-HCl, pH 8.3, 50 mM KCl, or LiCl for Li^+^ experiments, 1 mM MgCl_2_, 0.8 % glucose, 0.5 mg/ml BSA) in a total volume of 10 ul was incubated for 1h on the benchtop in the dark. Before addition of primer to the enzyme, a 10x primer stock was diluted in 1x T50 with LiCl and then heated for 5 min at 95°C. The 10x primer was snap cooled on ice and 1 ul was subsequently added to the enzyme. The imaging buffer was first saturated with Trolox (triplet state quencher) and then filtered through a 0.22 um filter and brought to pH 8.3. The enzyme/primer complex was then diluted with 40 ul of Ix imaging buffer and then incubated on the slide for 10 minutes. The slide was put on the TIRF objective, the density of immobilized molecules was assessed and 100 ul of imaging buffer + 1 ul gloxy solution (200 ug/ml catalase and 100 mg/ml glucose oxidase in T50) was flowed through the channel.

#### Primer extension by surface-immobilized telomerase

After the telomerase complexes were stalled on the slide surface, the indicated dNTPs were added as described in the text at a concentration of 200 uM. Reactions were allowed to proceed for the indicated amount of time. For real-time experiments, dNTPs were added to the slide during active data acquisition.

#### Data acquisition and analysis

Imaging fields containing 300-700 molecules were imaged at 2 frames/second for real-time smFRET experiments and 10 frames/second for standard smFRET experiments. Green laser intensity was set to 3mW for real-time experiments and to 15mW for standard experiments. 20 two-second movies were collected at each indicated time-point for the standard FRET experiments and 15-minute movies were collected for the real-time experiments. Individual traces were parsed out using custom written IDL software to correct for dye-cross talk and background. Individual traces were then filtered in MATLAB and molecules that did not contain and acceptor dye were discarded. FRET intensities were calculated using the equation I_A_/(I_A_+yI_D_) where I_A_ is the acceptor intensity, I_D_ is the donor intensity and y is the gamma correction factor. The first two seconds of individual FRET traces were binned into histograms.

### Kinetic analysis

For kinetic analysis, the primer extension assay described above was modified by chasing with 20 μM cold (TTAGGG)_3_ primer after 20 minutes of initiating the reaction, to prevent hot primer and hot product rebinding within the subsequent 70-minute time course. The intensities of the RAP bands on the gel electrophoretogram were converted to absolute concentrations based on the intensity of the initial hot primer band (50 nM). Multiple exposures were analyzed to ensure intense and weak bands remained within the linear range of the phosphorimager. These data were analyzed globally according to the sequential model (Figure 6B) using DynaFit (59, 60). Further details are given in the supplementary Methods.

## ACKNOWLEDGEMENTS

We thank Dr. Harry Noller and Dr. Alan Zahler for critical reading of the manuscript. This work was supported by National Institutes of Health Grants F99CA212439 (to L.I.J) and R01GM095850 (to M.D.S).

## AUTHOR CONTRIBUTIONS

L.I.J. and M.D.S. designed research, L.I.J., T.C. and R.B. performed research, L.I.J., J.H., J.W.P., C.R.B. and M.D.S. analyzed data and L.I.J. and M.D.S. wrote the paper.

## SUPPLEMENTARY METHODS

### Kinetic Analysis

The intensities of RAP bands resolved by gel electrophoresis were analyzed globally using DynaFit according to the mechanism shown in Figure 6A (1, 2). All rate constants were floated, except *k_a0_* and *k_d0_* whose values, within limits, had little effect on the analysis. The value of *k_a0_* is beyond the time resolution of the measurement (Fig. S6A) and was fixed at 1 nM^−1^ min^−1^ (with a lower limit of 0.1 nM^−1^ min^−1^), while *k_d0_* was fixed at ≤ 0.1 min^−1^. The association rate constant for the products to the telomerase enzyme was assumed to have the same *k_a0_* value as the primer. While this is unlikely to be strictly true, rebinding of products to the telomerase enzyme is likely to be slower, at least in the early stages of the reaction, when the hot primer remains in >5-fold excess over the sum of the dissociated products. Following a chase with 20 μM cold primer, binding of hot primer and labeled products is negligible and allowed for a more robust analysis without any assumptions regarding the precise values of *k_a_*. The cold chase data were fitted using the “incubate” command to model the preincubation of telomerase with the hot primer.

When modeling more than 10 sequential steps, DynaFit took many minutes to converge to a solution. The analysis could be accelerated by modeling the data in batches that overlapped by 2 repeats. For example, the estimated rate constants for the first 8 steps, obtained from fitting to the first batch of 10 steps, were set as fixed parameters in the analysis of the second batch of data which extended to 18 steps. This simplification is valid because *k_f_* is irreversible so that the steps are decoupled from each other. Batch wise analysis also allowed Monte-Carlo analysis to be performed on a practical time scale, to check the confidence of the fit and for any covariance between parameters (Fig. S5).

We also attempted to analyze time courses for primer extension over 90 minutes without a cold chase (Fig. S3) in which substrate depletion and product rebinding could be a complicating factor. The initial hot primer showed an initial rapid drop in concentration, followed by a slightly curved decay (Fig. S6A). The amplitude of the initial drop in primer concentration (5.3 nM) matched the initial burst in the sum of the first 3 dominant repeat bands (B, C and D) and provides a measure of the active telomerase concentration (3, 4). The small deviation of the primer decay profile from linearity indicates that the primer is depleted to an extent that *k*_*a0*_[primer] is no longer >> *k*_*d0*_ + *k*_*f0*_ and/or the DNA products are rebinding competitively with the primer. The effect is equivalent to deviation of the initial linear rate in classical steady-state enzyme kinetics due to substrate depletion and/or product inhibition. However, at the 20-minute time point, equivalent to the preincubation period in the cold chase experiment, the deviation is small (< 0.4 nM at a remaining primer concentration of 39 nM). Furthermore, when the complete time course of the first 12 repeats (B to M) were globally fitted using DynaFit, the saw-tooth pattern of the rate constants for sequential steps in the reaction remained (Fig. S6B).

The low intensity intervening bands corresponding to nucleotide addition processivity (NAP) were ignored in the analysis above. As a check, we analyzed a data set containing the five NAP bands between two RAP events and found their inclusion had little effect (< 10% change) on the rate constant estimates, *k*_*f*_ and *k*_*d*_ returned for the RAP bands. The intensity of the NAP bands were insufficient to extract any meaningful rate constants for nucleotide addition and dissociation.

**Figure S1.**
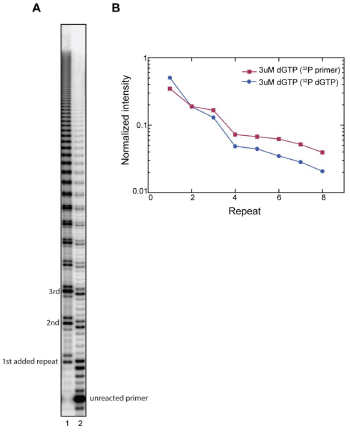
Using radiolabeled dGTP or radiolabeled primer result in similar product accumulation profiles. (**a**) Telomerase primer extension assay with 3uM dGTP and 500uM dATP and dTTP. Repeats added to the primer are indicated to the left. Lane 1 is performed with 0.3uM radiolabeled dGTP and 2.7uM cold dGTP. Lane 2 is performed with 50nM radiolabeled (TTAGGG)_3_ primer and 3uM cold dGTP. All assay conditions except for the identity of the radiolabeled nucleic acid are identical between the two samples. **b**) Normalized intensity plotted vs repeat number. Fractional intensity of each band compared to total lane counts is plotted against repeat number.

**Figure S2.**
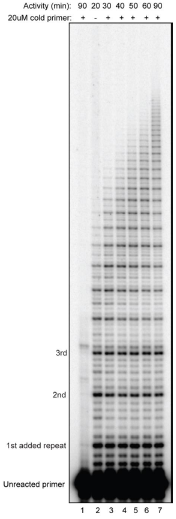
Telomerase is processive in the presence of 50nM primer. Telomerase primer extension assay with 20uM chase primer (TTAGGG)_3_ added after 20 minutes. Repeats added to the primer are indicated to the left.

**Figure S3.**
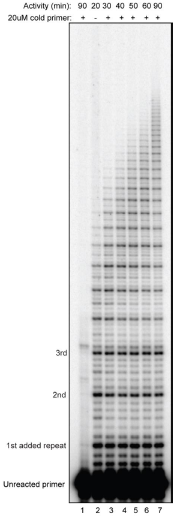
Telomerase activity is slower in Li^+^. Telomerase primer extension assay with samples taken at the indicated time points in either KCl (lanes 1-7) or LiCl (lanes 8-14). Repeats added to the primer are indicated to the left.

**Figure S4.**
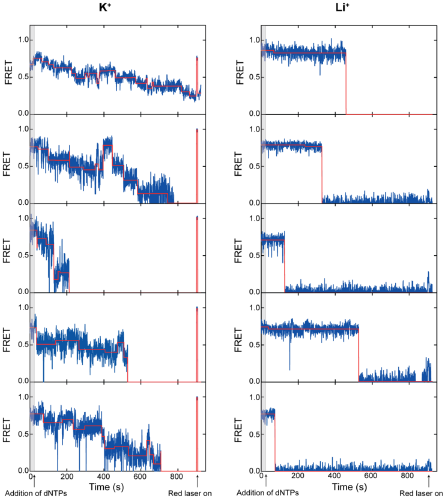
Real-time smFRET traces. Representative FRET traces are shown for experiments performed in KCl (left panels) and LiCl (right panels). Steps (red) were fit to each trace using Matlab. Red laser was turned on at 900 (s) to check for Cy5 photobleaching. dNTPs were added after 25 seconds (grey bar).

**Figure S5.**
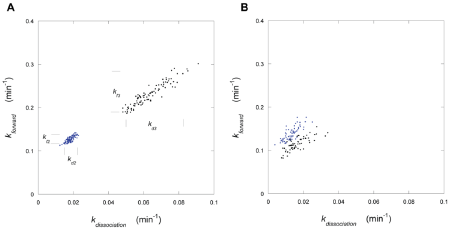
Example Monte-Carlo analysis to illustrate confidence of fitting and covariance of parameters for the scheme shown in Figure 6A. (1, 2). The plots show the rate constants for the dissociation and forward elongation for the repeat bands, C and D (Figure 6A) in the presence of (A) K^+^ and (B) Li^+^. The analysis shows that *k_f3_* is about 2-fold larger than *k_f2_* and *k_d3_* is about 3 times larger than *k_d2_* in the presence of K^+^, leading to reduced microscopic processivity (*k_f_*/(*k_f_*+*k_d_*)), whereas the differences in the presence of Li^+^ are marginally significant. The markers in (A) indicate the 5 and 95 percentile limits of the estimated rate constants based on over 500 iterations. This analysis indicates that the saw-tooth pattern of consecutive rate constants and the processivity observed in K^+^ is significant compared with the near-featureless pattern in Li^+^ (Figure 6C). The analysis also shows that *k_f_* and *k_d_* for each repeat band are linearly covariant, leading to an elliptical distribution. Consequently, the microscopic processivity is better defined than the individual rate constants. No covariance is seen between pairs of other rate constants (e.g. *k_f3_* versus *k_d2_*).

**Figure S6.**
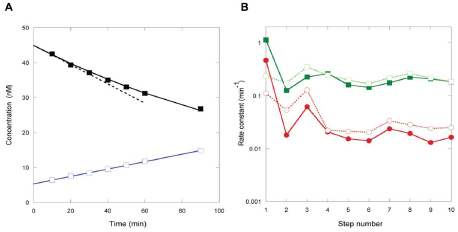
Reproducibility of kinetic analyses. (A) Primer extension assay performed in the absence of a cold chase, starting with 50 nM hot primer. Note the drop in primer concentration to 45 nM which is complete within the first time point (black solid symbols). This indicates that *k_on_* = *k*_*a0*_[primer] + *k*_*d0*_ + *k*_*f0*_ is rapid on the time scale of this experiment and is dominated by *k*_*a0*_[primer] based on modeling (Fig 6D; *k*_*f0*_ ≈ 0.1 min^−1^) and literature data (*k*_*d0*_ < 0.0006 min^−1^ (5). The solid black line shows an exponential fit to the data, and the dashed line shows the equivalent initial rate, to illustrate the small deviation from linearity of the early time points. The initial rapid drop in primer concentration is matched by a 5 nM burst in product formation, which is dominated by the first 3 repeat products (blue open symbols = sum B + B^#^ + C + C^#^ + D + D^#^). (B) The rate constants returned by global fitting using DynaFit for primer extension assays carried out in the presence of K^+^, with a 20 μM cold primer chase (solid symbols and line = experiment of Figure 6D) compared with a similar experiment (Fig. S3) in the absence of a cold chase (open symbols, dashed line) in which product rebinding may occur during the 90-minute incubation. These data indicate that the saw-tooth profile of rate constants (green squares = *k_f_* and red circles = *k_d_*) is a robust feature of telomerase that is observed using a different experimental design and a different sample preparation.

**Figure S7.**
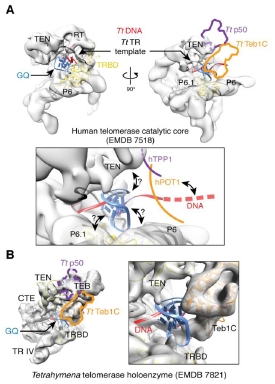
Model of a GQ on the telomerase enzyme. (A) The DNA exit channel in human telomerase provides space to accommodate a GQ. Structural superposition with the Tetrahymena telomerase EM structure places the hPOT1-hTPP1 counterparts TtTeb1-Ttp50 in juxtaposition to the DNA exit upstream of the GQ (outlines at right, see B). Bottom panel, close up of the GQ model. The distance of the GQ to the active site is below two telomeric repeats (12 nt) according to observations presented in this study. In this position, the GQ has access to critical elements such as the TEN domain, the TRBD and the hTR CR4/5 to exert its potential impact on telomerase function. Modeled coordinate PDB IDs: GQ, 2HY9; TR template and DNA, 6D6V; TRBD-P6-P6.1, 4O26. Human telomerase catalytic core EM reconstruction, EMDB 7518. (B) In Tetrahymena, the modeled GQ pocket is further confined by the C-terminal domain of the TEB complex protein Teb1, illustrating how telomere binding proteins can be excluded from a GQ promoting environment within telomerase holoenzyme complexes. Modeled coordinate PDB IDs: Tetrahymena TERT, TR, p50, Teb1 and DNA, 6D6V; GQ, 2HY9. Tetrahymena telomerase holoenzyme EM reconstruction, EMDB 7821.

**Table S1.**
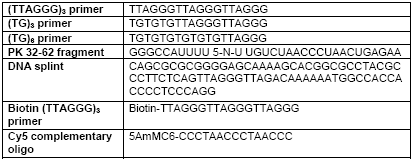
Oligos used for this study.

